# Probing the mechanisms of intron creation in a fast-evolving mite

**DOI:** 10.1101/051292

**Authors:** Scott William Roy

## Abstract

Available genomic sequences from diverse eukaryotes attest to creation of millions of spliceosomal introns throughout the course of evolution, however the question of how introns are created remains unresolved. Resolution of this question has been thwarted by the fact that many modern introns appear to be hundreds of millions of years old, obscuring the mechanisms by which they were initially created. As such, analysis of lineages undergoing rapid intron creation is crucial. Recently, Hoy et al. reported the genome of the predatory mite *Metaseiulus occidentalis*, revealing generally rapid molecular evolution including wholesale loss of ancestral introns and gain of new ones. I sought to test several potential mechanisms of intron creation. BLAST searches did not reveal patterns of similarity between intronic sequences from different sites or between intron sequences and non-intronic sequences, which would be predicted if introns are created by propagation of pre-existing intronic sequences or by transposable element insertion. To test for evidence that introns are created by any of multiple mechanisms that are expected to lead to duplication of sequences at the two splice boundaries of an intron, I compared introns likely to have been gained in the lineage leading to *M. occidentalis* and likely ancestral introns. These comparisons did initially reveal greater similarity between boundaries in *M. occidentalis*-specific introns, however this excess appeared to be largely or completely due to greater adherence of newer introns to the so-called protosplice site, and therefore may not provide strong evidence for particular intron gain mechanisms. The failure to find evidence for particular intron creation mechanisms could reflect the relatively old age of even these introns, intron creation by variants of tested mechanisms that do not leave a clear sequence signature, or by intron creation by unimagined mechanisms.

---

The ubiquity of spliceosomal introns in eukaryotic nuclear genes and the diversity of intron positions across eukaryotic diversity attests to a huge number of intron creation events in the history of eukaryotes (Rogozin et al. 2003). In 2003, Alexei Fedorov and myself performed near-genome-scale comparisons of intron positions in orthologs, in hopes of identifying recently-created introns (Roy et al. 2003). Much to our surprise (and chagrin), we found a remarkably small number of changes: among 10,020 intron positions studied in a comparison of species diverged 80 million years ago (human and mouse), we found only five that were not shared between the species. Furthermore, all five of these were shared with outgroups, indicating intron loss and not gain (Roy et al. 2003). Other genome-wide and smaller-scale studies have confirmed the general finding of striking degrees of intron position sharing suggestive of little intron gain in many different lineages over a variety of phylogenetic depths (Stajich and Dietrich 2005; Roy and Hartl 2006; Roy et al. 2006; Rogozin et al. 2003; Yang et al. 2013).

This finding of small numbers of intron gains in many lineages has greatly hindered our understanding of the mechanisms, phenotypic impacts and population genetics of intron-exon structures. However, a growing handful of studies have begun to fill in these long-standing gaps. Li et al. (2009, 2014) and Omilian et al. (2008) discovered dozens of recent intron creations in the water flea *Daphnia pulex* which appear to have arisen by imprecise double strand break repair (DSBR), and Farlow et al. (2011), Yenerall et al. (2011) and Sun et al. (2014) have provided evidence that a similar mechanism may create introns in species of *Drosophila* and *Neurospora*. Alex Wordens group and others have probed widespread creation of introns in the green alga *Micromonas pusilla* by propagation of a cryptic transposable element (Worden et al. 2009; van Baren et al. 2016; Simmons et al. 2016; Verhelst et al. 2013), and multiple groups have reported on a seemingly similar case in a clade of fungi (Collemare et al. 2013, 2015; van der Burgt et al. 2012). Curtis and Achibald (2010) reported a single intron gained by insertion of a non-intronic portion of mitochondrial DNA. We previously reported a variety of intron gain mechanisms in the fast-evolving chordate *Oikopleura*, including local propagation of short intron sequences by unknown mechanisms and intron creation by transposable element insertion (Denoeud et al. 2010). We also previously reported creation of introns by intronization, i.e. splicing out of internal portions of ancestral exonic sequences (Irimia et al. 2007). Hellsten et al. 2011 reported intron creation by internal duplication of exonic sequencing and usage of two resulting copies of an AG|GT containing motif (Hellsten et al. 2011).

However, even while revealing much about intron creation, the diversity of mechanisms reported by these studies increases the importance of finding more cases, in order to understand the relative incidence and determinants of these different mechanisms across species. Here, I study the recently-reported genome of the predatory mite *M. occidentalis*, which has undergone widespread intron gain over long evolutionary timescales (Hoy et al. 2016).

## Sequence similarity searches do not support intron creation by intron propagation or transposable element insertion

To test for the possibility of insertions of introns at new sites by propagation of pre-existing introns to new sites, I used BLAST comparisons between regions including intronic sequences and their flanking exonic sequence. The expected signature of intron propagation would be sequence similarity including the entire intron sequence, but not flanking exonic sequences. I downloaded the *M. occidentalis* genome and annotations from Genbank (GCF_000255335.1_Mocc_1.0), and extracted every annotated intronic sequence within the *M. occidentalis* genome along with 20 exonic nucleotides on either side (for 52,196 introns in total). I performed all-against-all BLASTN searches (with filtering of repetitive sequences turned off in order to promote extensions of BLAST hits through low-complexity regions) between these sequences, and identified hits that began and ended within 10 nucleotide positions of the splice boundaries for both the query and subject sequences. This identified only a single case (a pair of introns in genes XP_003744432.1 and XP_003744441.1), which upon manual alignment was revealed to involve longer homologous regions extending far beyond the ends the introns (Figure 1a). Thus this search revealed no case of inter-intron similarity that would be suggestive of intron propagation.

**Figure 1.**
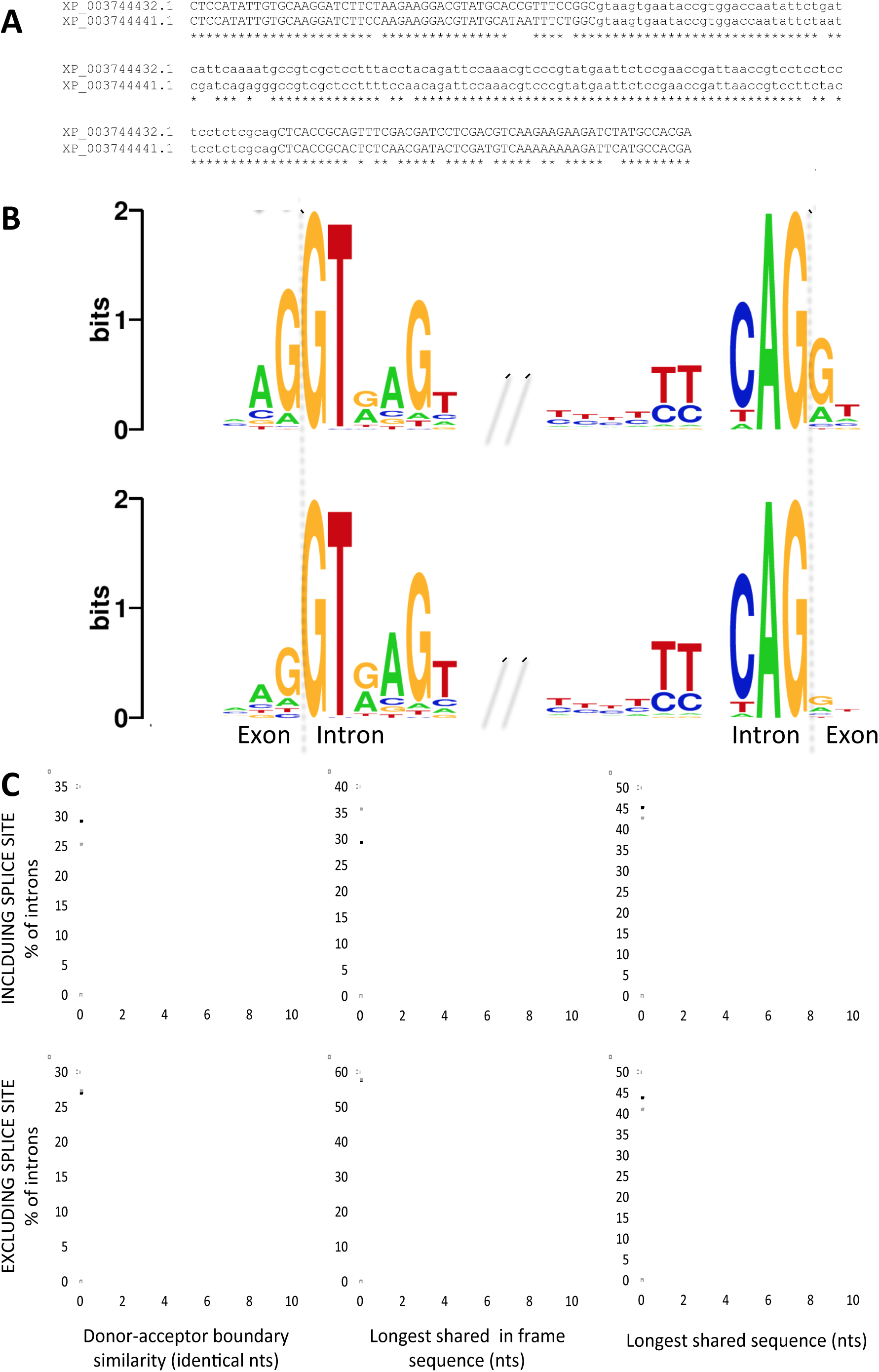
Search for signatures of intron creation mechanism in *M. occidentalis* reveals no clear patterns. A. The single pair of introns for which initial screening for signatures of intron propagation instead represents an extended region of homology. Clustalw2 alignment of similar introns with flanking exons within genes XP_003744432.1 and XP_003744441.1 are shown. Upper/lowercase indicates exonic/intronic nucleotides. B. Sequence logo plots for novel (top) and shared (bottom) introns, showing greater protosplice character at the two flanking nucleotide sites of each exon. C. Comparison of sequence similarity between donor and acceptor sites for novel (black) and shared (gray) introns. Left: number of nucleotide identities within ungapped alignment of 10 nucleotides spanning donor and acceptor boundaries for each intron. Middle: Longest number of shared nucleotides within ungapped alignment of 10 nucleotides spanning donor and acceptor boundaries for each intron. Right: Longest total shared motif between extended donor and acceptor regions (including 5 exonic and 15 intronic nucleotides). Results are given either including (top) or excluding (bottom) 2 terminal nucleotide positions for each exon/intron.

I next tested for evidence of intron creation by insertion of another genomic sequence (in particular a transposable element, although the case reported by Curtis and Archibald (2010) raises the possibility of a copy of a non-mobile sequence being inserted into a new genomic locus). Again, the expected sequence signature in this case would be a region sequence similarity corresponding closely to the boundaries of the intron. I performed a BLASTN search of all of the sequences generated above (intron plus 20 nucleotides of flanking exonic sequence) against the entire *M. occidentalis* genome and, as above, identified hits that began and ended within 10 nucleotide positions of both the query and subject sequences. This identified a total of 10 introns with genomic hits (several with multiple hits). As with the intron-intron case, for each of the 10 cases, manual scrutiny and alignment revealed multiple genomic copies extending well into the annotated exonic sequences, indicating that these cases do not reflect intron creation from a genomic sequence newly inserted into the interior of an exon. In addition, specific BLASTN searches against the mitochondrial genome revealed no similarity to nuclear intronic sequences.

## Novel introns have stronger protosplice sites but do not have extended shared motifs at their boundaries

I next tested a prediction of several different hypotheses, namely similarities between the sequences spanning or flanking the two splice sites (donor and acceptor). Such similarities are expected by multiple models including DSBR, in cases involving sticky end breakage and repair (Li et al. 2009), and internal duplication of exonic sequencing and usage of two resulting copies of an AG|GY containing motif (Rogers 1989; Venkatesh et al. 2009; Hellsten et al. 2011). Testing for similarity between the two boundaries of an intron is complicated by the lack of a clear null expectation, particularly given that introns in general in several species (including ancestral introns) exhibit sequence preferences at the termini of the flanking exons that match the corresponding intronic sequence (AG|gt donor and ag|GT acceptor).

This complication can be circumvented by specifically testing for an excess of donor-acceptor similarity in putatively more recently created introns relative to putatively more ancestral introns, since the former are more likely to retain sequence signatures betraying their mechanism of creation. I compared intron positions in putatively orthologous genes between (i) *M. occidentalis*, (ii) the tick *Ixodes scapularis*, the closest relative for which a genome is available; and (iii) *Homo sapiens*, chosen because of its retention of a large fraction of the ancestral metazoan intron complement (see e.g., Srivastava et al. 2008). The *I. oxodes* and *H. sapiens* genomes and annotations were downloaded from Genbank, intron-exon structures were extracted, and orthologs were defined by reciprocal BLAST searches at the protein level. Genes were aligned at the protein level in Clustalw2 with standard parameters, and conserved protein-coding regions identified were defined as positions with ≥40% amino acid identity and no gaps for windows of 10 amino acid positions on both sides of a position, for all three pairs of species individually (use of a variety of less stringent criteria did not qualitatively change any of the following results; data not shown). Intron positions were then mapped onto the protein-level alignment and each *M. occidentalis* intron position was identified as novel or as shared (i.e., an intron position at which an intron in *I. scapularis* and/or *H. sapiens* is found at the exact homologous position in the alignment), as in Rogozin et al. (2003) or Roy et al. (2003). This identified 3355 novel and 2509 shared *M. occidentalis* introns. Notwithstanding the ongoing debate about whether shared intron positions reflect actual shared ancestral introns (Li et al. 2014), for brevity these will be referred to as shared/novel introns (in place of the more precise introns at shared/novel positions’).

Figure 1b shows overall sequence logos of splice boundaries for the two sets of introns (Crooks et al. 2004). Interestingly, this comparison reveals a clearly much stronger preference for the so-called protosplice site in new introns (*P* < 10^−10^ for each of the −1, −2 and +1 bases individual, by simulation). This finding is consistent with previous results (Sverdlov et al. 2003; Qiu et al. 2004); however, inference of intron age in the previously analyzed datasets was a notoriously difficult endeavor (Rogozin et al. 2003; Roy and Gilbert 2005; Csuros 2005; Carmel et al. 2007), thus observation of the same pattern on this simpler dataset is a comforting confirmation.

To test for similarity between donor and acceptor sites of the same intron, I performed three similar but distinct tests. First, for each intron I performed an ungapped alignment of the 10 nucleotides straddling the donor and acceptor boundaries (5 nucleotides on either side of the splice boundary) and counted the number of nucleotide matches. Second, I counted the maximum number of such matches in a row within this same region. Third, I compared the maximum length of shared motif between donor and acceptor site regions within an extended 20 nucleotide region (five exonic nucleotides plus 15 intronic nucleotides from donor and acceptor). All three comparisons did show differences in the predicted direction, with clearly more matches, clearly longer runs of matches, and very slightly more overall longer shared motifs in novel than shared introns (top row of Figure 1c, left, middle and right, respectively; *P* < 0.01 for each test). However, because the AG|G protosplice site matches the corresponding intron positions (AG|g…ag|G), it is possible that this excess simply reflects greater similarity at the boundary due to the strong protosplice site in novel introns (which may or may not reflect the mechanism of intron creation; see below). Therefore I ran equivalent tests excluding the four positions flanking the splice sites (the NNgt and agNN sites; thus positions −7 through −3 and +3 through 7 were compared for total matches and run of matches, 7-to-3 for exonic side and 17-to-3 for intronic side for longest total shared motif). These tests showed a very different pattern (bottom row of Figure 1c). All three tests showed no clear difference between novel and shared introns: thus there is no evidence for greater similarity between 5’ and 3’ boundaries for novel introns outside of the stronger protosplice site.

## Concluding remarks

Examination of intronic sequences in a metazoan with highly divergent intron-exon structures for the most part failed to reveal expected signatures of several tested mechanisms. The one clear difference between novel and shared introns was the greater strength of the protosplice site. The implications of this finding are not clear. Greater protosplice site character is exactly as expected from a variety of mechanisms that lead to duplication of the exonic insertion site; however, greater protosplice character could also reflect greater success of newly-created introns that insert into optimal exonic contexts (e.g., the upstream exonic AG can participate in basepairing to the U1 snRNA, a key step in spicing).

The lack of clear sequence signatures of intron creation mechanisms is also difficult to interpret. While the divergent character of the *M. occidentalis* genome relative to other available metazoan genomes attests to a large degree of change, no genomic sequences are available for close relatives of *M. occidentalis–I. scapularis*, the closest relative with a genome sequence, is from a different order – thus the timing of these changes is unknown. Most of the intron creation in the history of *M. occidentalis* could date to many million of years ago, in which case it would not be surprising that the sequence signatures of intron creation are no longer detectable. Alternatively, the introns in *M. occidentalis* could have been gained by mechanisms without clear sequence signatures. For instance, in the data of Li et al. (2009, 2014) some newly–gained introns do not show clear tandem repeats at the intron boundaries, which could reflect intron creation by repair of a blunt end double strand break (which is not expected to result in tandem duplication of a sequence at the borders). Still another possibility is suggested by the extremely rapid sequence evolution of new introns in *D. pulex* discovered also reported by Li et al. (2014), which could rapidly obscure the origins of new introns. Finally, introns in *M. occidentalis* could be created by a mechanism not yet imagined (or at least not tested here). Genomic sequences from closer relatives of *M. occidentalis* will hope to clarify the evolutionary history of this intriguing transformed genome.

